# Cryo-EM structures of a bathy phytochrome histidine kinase reveal a unique light-dependent activation mechanism

**DOI:** 10.1101/2024.05.17.594632

**Authors:** Szabolcs Bódizs, Petra Mészáros, Lukas Grunewald, Heikki Takala, Sebastian Westenhoff

## Abstract

Phytochromes are photoreceptor proteins in plants, fungi and bacteria. They can adopt two photochromic states with differential biochemical responses. The structural changes transducing the signal from the chromophore to the biochemical output modules are poorly understood due to challenges in capturing structures of the dynamic, full-length protein. Here, we present the first cryo-electron microscopy structures of the phytochrome from *Pseudomonas aeruginosa* (*Pa*BphP) in its resting Pfr and photoactivated Pr state. The kinase-active Pr state has an asymmetric, dimeric structure, whereas the kinase-inactive Pfr state opens up. This behaviour is different from other known phytochromes and we explain it with the unusually short connection between the photosensory and output modules. Multiple sequence alignment of this region suggests evolutionary optimisation for different modes of signal transduction in sensor proteins. The results establish a new mechanism for light-sensing by phytochrome histidine kinases and provide input for the design of optogenetic phytochrome variants.

## Introduction

Life depends on light, and living organisms have evolved sophisticated systems to sense it and adjust accordingly. Phytochrome proteins enable red/far-red photoreception by converting the captured photosignal into structural changes translating to biochemical activity. In the 1950s, researchers discovered phytochromes in plants as they tried to understand how plants control periodic events, such as flowering^1^. Since then, phytochromes have been identified in many different organisms including fungi, cyanobacteria, and even non-photosynthetic bacteria^2,3^. These organisms all utilise phytochrome photosensation to control essential biological and light-dependent processes like growth, reproduction, and movement^4^.

Phytochromes function by interconverting between two distinct photochromic states: the Pr state that absorbs red light with a peak absorption at 700 nm, and the Pfr state that absorbs far-red light with a peak excitation at 750 nm. These two structurally different states have different signalling activities. In natural lighting, the protein exists in an equilibrium between the two states, with shifts in this equilibrium depending on the illumination conditions^4^. The structural photoconversion process between the two states and the connection of the photoequilibrium to biochemical function has raised considerable scientific interest^5–14^. Two types of phytochromes are defined based on the relative free energies of the two states: *canonical* phytochromes have a more stable Pr state, whereas *reverse-activated*, or *bathy* phytochromes have Pfr as their resting state. A deeper understanding of the photoactivation process can be applied in the design of near-infrared fluorescent markers in microscopy or optogenetic tools, where phytochrome constructs are readily used as sensory modules^15–18^.

Phytochromes have a homodimeric modular structure, typically composed of an N-terminal photosensory module and a C-terminal effector/output module (Fig. 1A)^19^. The photosensory module is conserved across the protein superfamily, consists of PAS (Per/Arnt/Sim), GAF (cGMP phosphodiesterase/Adenylyl cyclase/Fhl1), and PHY (PHYtochrome-specific) domains, and contains a bilin cofactor as the chromophore. A flexible loop of the PHY domain, called the PHY-tongue, undergoes considerable structural changes upon photoactivation^5^. The configuration of the output module is more variable: in many bacterial phytochromes, it features a HK (histidine kinase) unit consisting of a DHp (Dimerisation/Histidine phosphotransfer) and a CA (Catalytic ATP-binding) domain.

**Figure 1.**
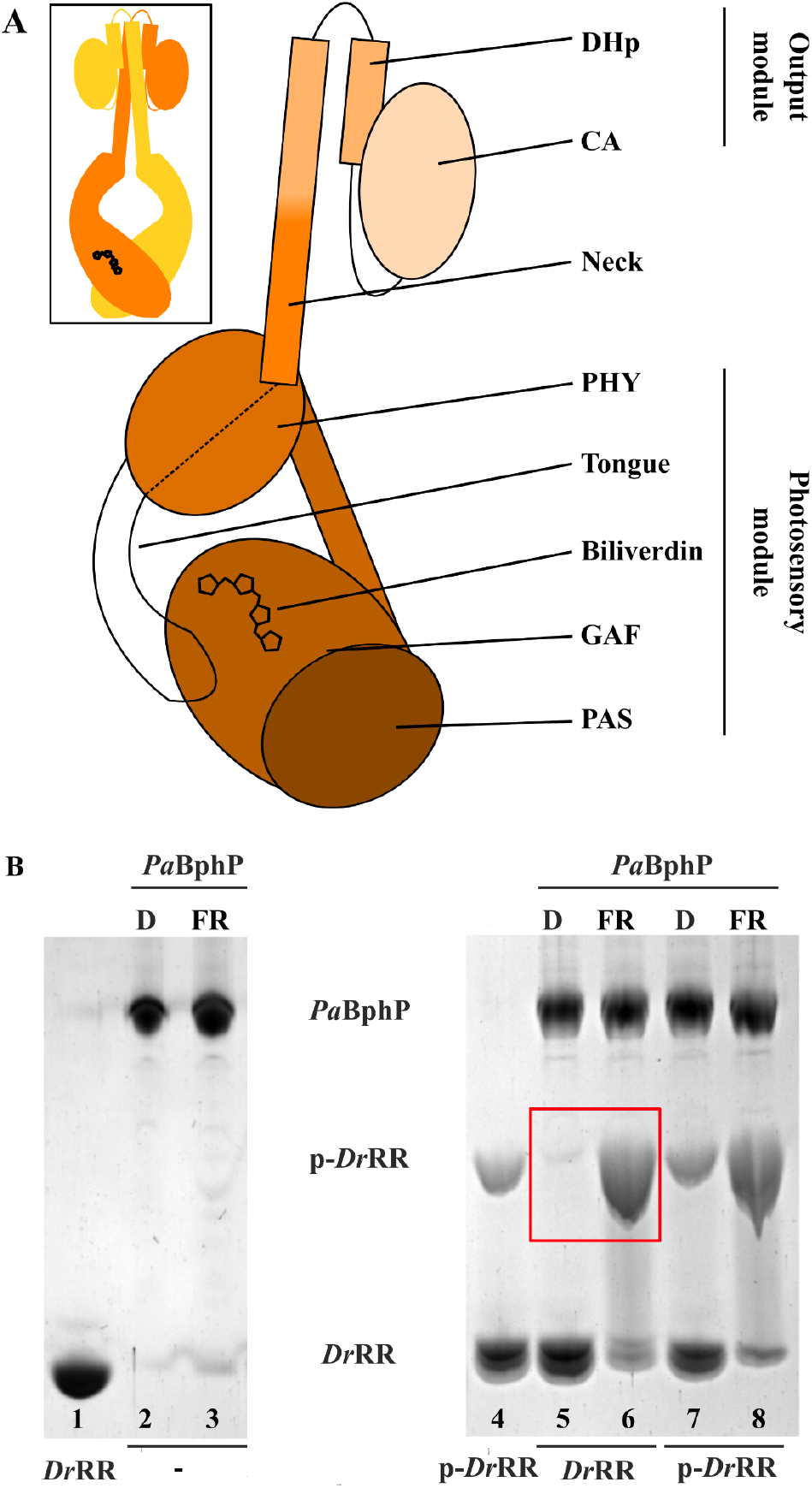
Schematic representation of bacterial phytochromes and the enzymatic activity of *Pa*BphP. **A** The head-to-head homodimeric configuration of the functional protein (inset), and the domain structure of a single chain. Domain abbreviations: CA (Catalytic ATP-binding), DHp (Dimerisation/Histidine phosphotransfer), GAF (cGMP phosphodiesterase/Adenylyl cyclase/Fhl1), PAS (Per/Arnt/Sim), PHY (PHYtochrome-specific). **B** The enzymatic activity of *Pa*BphP, determined with Phos-tag assay, which resolves phosphorylated proteins from non-phosphorylated ones in a gel^12^. The migration of the phosphorylated response regulator from *D. radiodurans* (p-*Dr*RR) is slower than its non-phosphorylated counterpart (*Dr*RR). The samples were supplemented with identical amounts of either *Dr*RR or p-*Dr*RR, and the change in the p-*Dr*RR content speaks for net kinase (increase) or phosphatase (decrease) activity. The assay indicates that *Pa*BphP phosphorylates *Dr*RR under far-red illumination (FR) but not in the dark (D). No phosphatase activity (i.e., disappearance of p-*Dr*RR input) was detected. See Figure S2 for an extended gel.

The first phytochrome crystal structure, comprising the PAS-GAF fragment of the *Deinococcus radiodurans* phytochrome, was obtained in 2005^20^. Subsequently, structures of the phytochrome sensory modules were acquired from Cph1 (*Syneochocystis sp*. PCC6803) and *Pa*BphP (*Pseudomonas aeruginosa*)^8,21^, with structures from various other species emerging later^5,22–29^. More recently, cryo-electron microscopy (cryo-EM) has provided structures of plant and bacterial phytochromes with output modules^30–33^. This approach has yielded insight into the structure of phytochromes in their native full-length, dimeric forms. However, there is a scarcity of structures for bathy phytochromes and those in photoactivated states, limiting progress in understanding the photoactivation mechanism in the context of full-length phytochromes.

HK containing phytochromes are part of a two-component signalling system. Typical two-component systems consist of transmembrane sensor kinases and response regulator proteins^34^. The sensor kinases are homodimeric proteins, which detect external signals and transduce these through the membrane, thereby modulating its intracellular kinase activity^35–38^. Due to the presence of hydrophobic regions that pass the lipid bilayer, obtaining comprehensive structural information for transmembrane HKs remains challenging^39–41^. As light can pass freely through the membrane, phytochrome kinases are soluble, intracellular proteins, but have a similar structural arrangement and interdomain communication to transmembrane HKs. This makes them valuable models for investigating the signal perception and transduction mechanism in two-component systems.

An important structural element shared between transmembrane kinases and phytochrome kinases is a coiled-coil alpha helix connecting the sensor and HK modules, which we refer to as the ‘neck’ (Fig. 1A). This neck precedes the catalytic histidine of the DHp domain and plays a role in signal transduction, serving as the sole connection between the input and output modules. The neck region, defined here as spanning from the PHY domain to the catalytic histidine (H513 in *Pa*BphP), appears to be important in this signal transduction process, but its mechanism of switching between states is poorly understood.

Depending on the photochromic state, bacterial phytochromes can function as a net kinase or phosphatase. In general, phytochrome histidine kinases show kinase activity in the Pr state, whereas some phytochromes act as a phosphatase in the Pfr state^9,11,12,42^. In particular, autophosphorylation assays of the bathy phytochrome *Pa*BphP have revealed reduced kinase activity in the dark, which suggests that bathy and canonical phytochromes have a comparable activity pattern^11,42^. In cellular context the phosphate signal has to be transduced to the response regulator protein. In vitro assays targeting the phosphotransfer reaction have only been reported for some bacterial phytochromes^12,42–45^. Here, we report a characterisation of this process for the bathy phytochrome *Pa*BphP.

In this study, we use single-particle cryo-EM to reveal the structure of the full-length phytochrome from *Pseudomonas aeruginosa* (*Pa*BphP) in both its resting (Pfr) and photoactivated (Pr) state. We provide detailed structural models for the protein in both states, establishing how photoactivation works in bathy phytochromes. We identify a new mode of photo(de)activation of the output domains, and suggest that this is controlled by the constitution of the neck region. Multiple sequence analysis on a broad range of HK-containing phytochromes highlights a potential evolutionary drive for distinct activation mechanisms.

## Results

### PaBphP is a bathy phytochrome with kinase activity in Pr

Our UV/VIS spectra of *Pa*BphP confirm the bathy character of the phytochrome as it relaxes towards the Pfr state in darkness (Fig. S1A)^8,46^. Generally, spectra of phytochromes reflect a photo-equilibrium between the Pr and Pfr states. For canonical phytochromes, the Pfr state can never be obtained in a pure form, because the Pr state cannot be selectively photoexcited. For bathy phytochromes, dark reversion leads to accumulation of Pfr. However, even in this case, a small percentage (<10%) of the proteins is expected to remain in Pr due to thermal activation^29,47^. Nevertheless, using a bathy phytochrome provides the opportunity to obtain an almost pure Pfr state for single particle cryo-EM investigation.

To establish enzymatic activity of *Pa*BphP, we conducted a Phos-tag phosphorylation assay (Fig. 1B). The assay probes if the phosphate is transduced from the phytochrome to a response regulator protein, which is the step that follows autophosphorylation in two-component signalling cascades. To establish this, we incubated *Pa*BphP with a phytochrome response regulator (RR) and ATP in the dark and under far-red (785-nm) illumination. As the cognate response regulator of *Pa*BphP is not known, we used the response regulator from *Deinococcus radiodurans* (*Dr*RR) as a substitute, which has been shown to be the phosphotransfer target of other phytochrome histidine kinases^12^. Indeed, we found out that *Pa*BphP can phosphorylate *Dr*RR in a light-controlled fashion. The p-*Dr*RR fraction is enriched under far-red illumination when supplying *Dr*RR and p-*Dr*RR as a phosphorylation target (lanes 6 and 8). Moreover, the net amount of p-*Dr*RR appears unaffected by the presence of *Pa*BphP in dark (lanes 5 and 7). This indicates that *Pa*BphP acts as a net kinase and transduces the phosphate signal in Pr, but is inactive in Pfr. Compared to other bacteriophytochromes, the on/off ratio appears high^12,48,49^.

### Single-particle cryo-EM structure of full-length PaBphP reveals a closed, asymmetric dimer

To link the functional changes to structure, we conducted cryo-EM on *Pa*BphP specimens prepared in the dark and under saturating far-red illumination. The illumination was supplied immediately prior to grid freezing, maximising the concentration of Pr in these grids (Fig. S1). We therefore expected that over 90% of the particles adopt Pfr in dark and Pr under far-red illumination. Single-particle cryo-EM datasets from both types were used to reconstruct near-atomic resolution structures. In the far-red illuminated (Pr) dataset, the final EM map (Fig. 2A) features reliable densities across the entire length of the protein, allowing us to build an atomic model (Fig. 2B). In the dark (Pfr) dataset, only the photosensory module (PAS-GAF-PHY) was resolved (Fig. 4).

**Figure 2.**
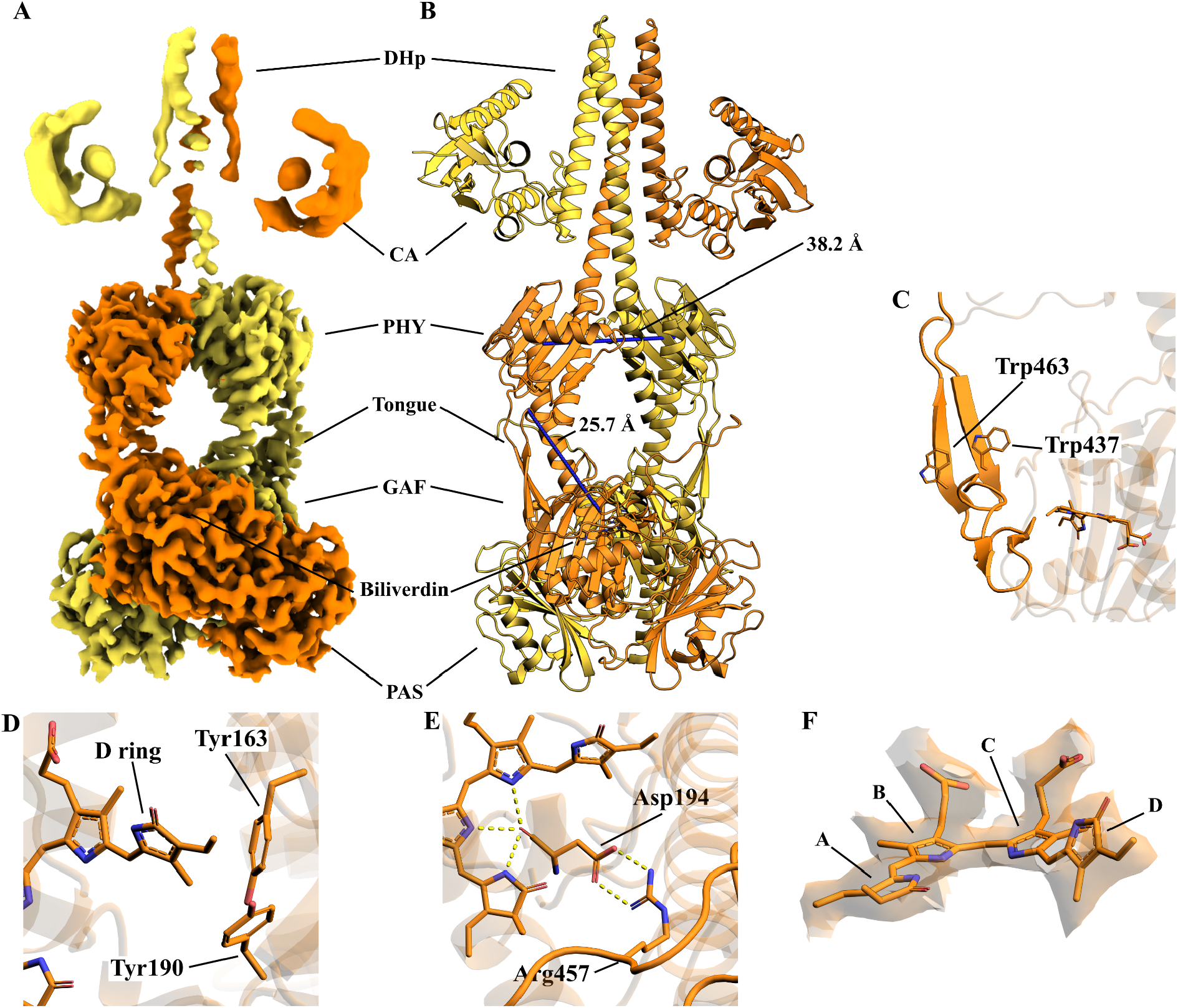
Structure of *Pa*BphP in its Pr state. **A** Composite EM density map (EMD-19981) showing the full-length protein. The chains are coloured differently to show coiling of the two protomers around each other. **B** Atomic model (9EUT) built into the density map. The PHY-tongue is extended, and the PHY domains make contact with each other. **C** The PHY-tongue is in a β-sheet conformation, with W437 anchoring it to the GAF domain. **D** Rotamerisation state of two tyrosine residues stabilising the conformation of the biliverdin D ring. **E** Polar contacts between the biliverdin, the PASDIP motif, and the PRxSF motif. **F** The density around the biliverdin, supporting the 15Z conformation.

We first describe the Pr structure obtained under far-red illumination (Fig. 2). In the collected dataset, 889,461 single particle images contribute to a highly homogeneous density of the photosensory module. However, there is considerable conformational heterogeneity in the output module region (Movie S1), which necessitated separate local refinements of the photosensory core and output module regions (Fig. S3). The local refinements were combined into a composite map (Fig. 2A), which was used to reconstruct the structure (Fig. 2B).

The density in the photosensory core region was reconstructed at a resolution of 3.0 Å, with densities for the chromophore and most side chains clearly visible. We find that the chromophore is in the 15Z conformation, which is expected for the Pr state (Fig. 2F). The D ring is positioned near the tyrosine residues Y163 and Y190, which are in typical orientation for Pr (Fig. 2D)^8,50^. Moreover, D194 of the conserved PASDIP motif is within hydrogen bonding distance of R457 of the PRxSF motif, establishing contact to the PHY-tongue (Fig. 2E). As expected for phytochromes in the Pr state, the tongue loop is in a β-sheet conformation (Fig 3C), with W437 making contact with the surface of the GAF domain^5,23^. The distance between D194 and G469, indicative of the length of the tongue, is 25.7 Å (Fig. 2B). Interestingly, we did not find clear electron density at the position of the pyrrole water (Fig. S4). The reason for this remains unclear. Even though one ordered water was visible at the acquired resolution in the electron density map (Fig. S4) and the pyrrole water is typically found to have low B-factors in crystal structures, the resolution of the map is borderline with some residues showing blurred densities and no other waters in the binding pocket observed.

**Figure 3.**
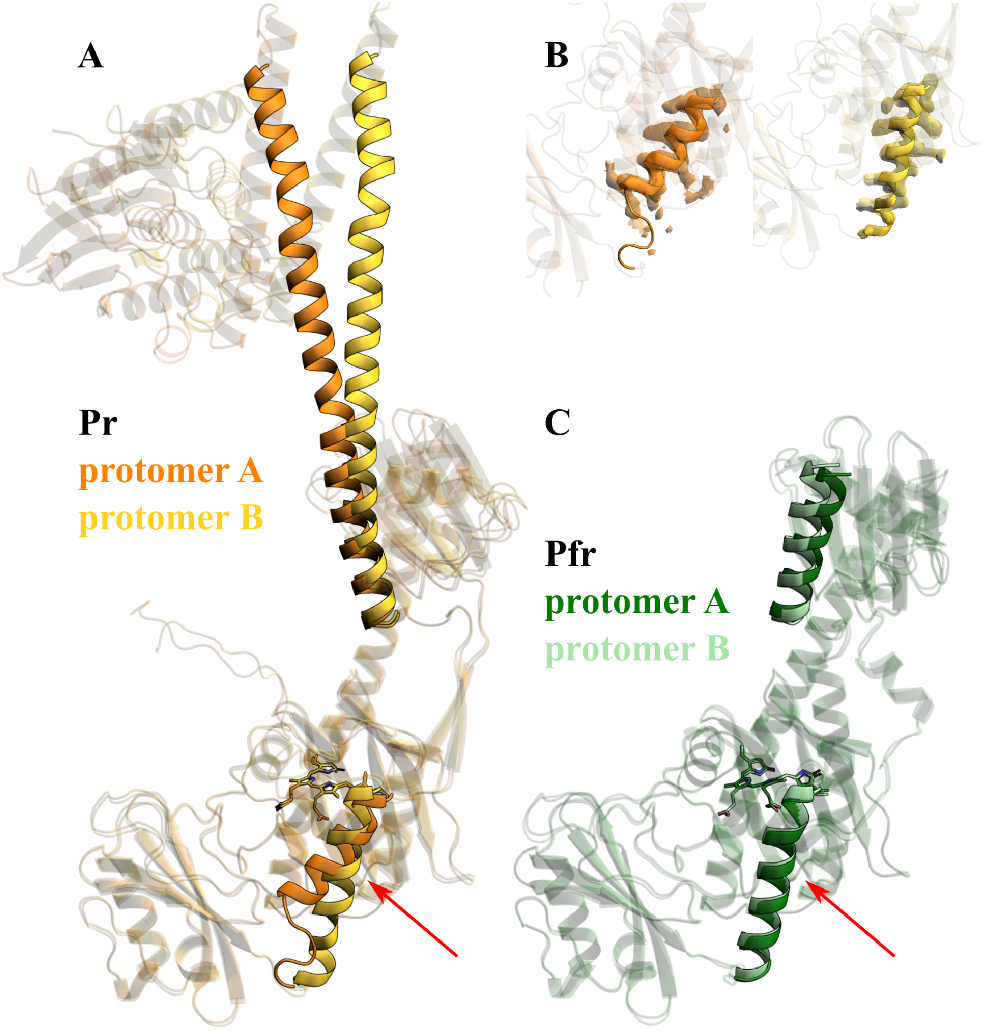
Symmetry in the structures of the Pr and Pfr states. The two protomers from each state were overlaid to reveal any symmetry breaking features. **A** The Pr structure is asymmetric in the PAS/GAF region, with helix A (red arrow) and the neck of protomer A tilting towards protomer B. **B** Density maps around helix A in protomers A/B in the Pr state support the asymmetry in the photosensory module. **C** The Pfr structure is symmetric in the entire photosensory core module, with helix A assuming the orientation observed in crystal structures.

The density of the output module was refined separately, with an estimated resolution of ∼10 Å. The map shows clear densities for the DHp and the N box helix of the CA domain. The peptide backbone of the DHp domain can be traced through the density, allowing the determination of the degree of coiling and revealing the direction of the hairpin connecting the four helix bundle. While the CA domain has lower interpretability, the N box helix density allows the placement of a predicted structure of the domain in the envelope of the density with high confidence in the position and orientation. The full-length model of *Pa*BphP in its Pr state is shown in Fig. 2B.

The structure is similar to two other structures in Pr, obtained from prototypical bacteriophytochromes^8,50^. This establishes that the Pr state of a phytochrome is similar, irrespective if the protein has Pr or Pfr as resting state. In contrast to the previous structures, we can position the DHp and CA domains with confidence. Importantly, the connecting loops between the DHp domain helices were fully resolved^51^. The loops reveal that the phosphorylation reaction will occur in between the CA domain of protomer A and catalytic histidine of protomer B (and *vice versa*), which is known as *trans*-autophosphorylation (Fig. S5)^30,31^. The orientation of the domains suggests a kinase-active state^52^.

The overall shape of the Pr dimer is asymmetric, with protomer A bent around protomer B at both dimerisation interfaces (Fig. 3A). While helix A (residues 118-136) of the GAF domain in protomer B assumes the orientation observed in crystal structures, the same helix in protomer A is tilted outwards at an angle of ∼13 degrees (Fig. 3B)^8^. Complementing this, the neck helix of protomer A is bent significantly, resulting in the HK dimer tilting towards protomer B. Asymmetry in the dimer arrangement has been observed in plant phytochromes, but not bacterial phytochromes^33^. It may suggest a tension in the Pr state structure.

### The structure of full-length PaBphP in Pfr reveals an open dimer

Now, we turn our attention to the dataset recorded in the dark. Here, two clearly distinct structures of the protein can be observed (Fig. S6). A small fraction of particles (approximately 10%) show a closed conformation (marked as “closed” in Fig S6). Since the refined Pr structure overlays very well with that volume, we assign these particles to residual Pr state. The majority (90%) of the particles show a different conformation, with features expected from Pfr in the photosensory module^8^. In this state, the chromophore is in the 15E configuration (Fig. 4F) and the tongue loop is rearranged to an α-helical conformation (Fig. 4C). In this α-helical form, W437 is positioned on the surface of the protein, and W463 takes its place in anchoring the tongue to the GAF domain^23^. The PASDIP-PRxSF connection is also rearranged compared to Pr, with D194 in contact with R453 and S459 (Fig. 4E). The PHY-tongue, as measured by the D194-G469 distance (20.1 Å), is 6 Å shorter than in Pr (Fig. 4B). Notably, the N-terminal strand of the loop folds into a second, shorter helix, which has not been observed in crystal structures of *Pa*BphP (Fig. 5), but is present in the crystal structures of bathy phytochromes Agp2 (6G1Y) and *Rp*BphP1 (4GW9)^8,26,53^.

**Figure 4.**
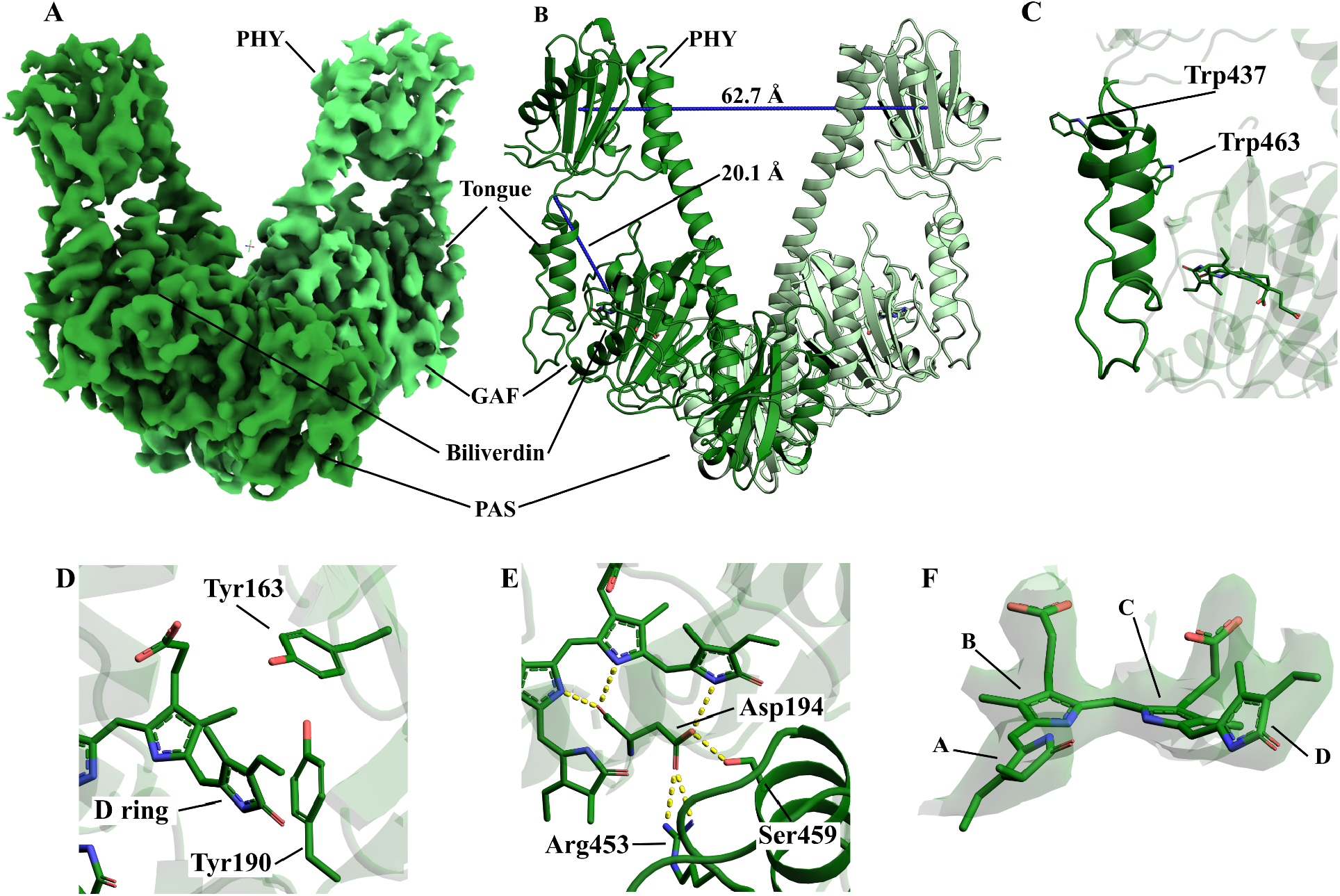
Structure of *Pa*BphP in its Pfr state. **A** EM density map (EMD-19989) showing the photosensory module of the protein. The chains are coloured differently to show the relative position of the two protomers. Flexibility of the DHp domain leads to complete loss of signal from the output module. **B** Atomic model (9EUY) built into the density map. The PHY-tongue is contracted, and the PHY domains are separated. **C** The PHY-tongue is in an α helical conformation, with W463 anchoring it to the GAF domain. **D** Rotamerisation state of two tyrosine residues stabilising the conformation of the biliverdin D ring. **E** Polar contacts between the biliverdin, the PASDIP motif, and the PRxSF motif. **F** The density around the biliverdin, supporting the 15E conformation.

**Figure 5.**
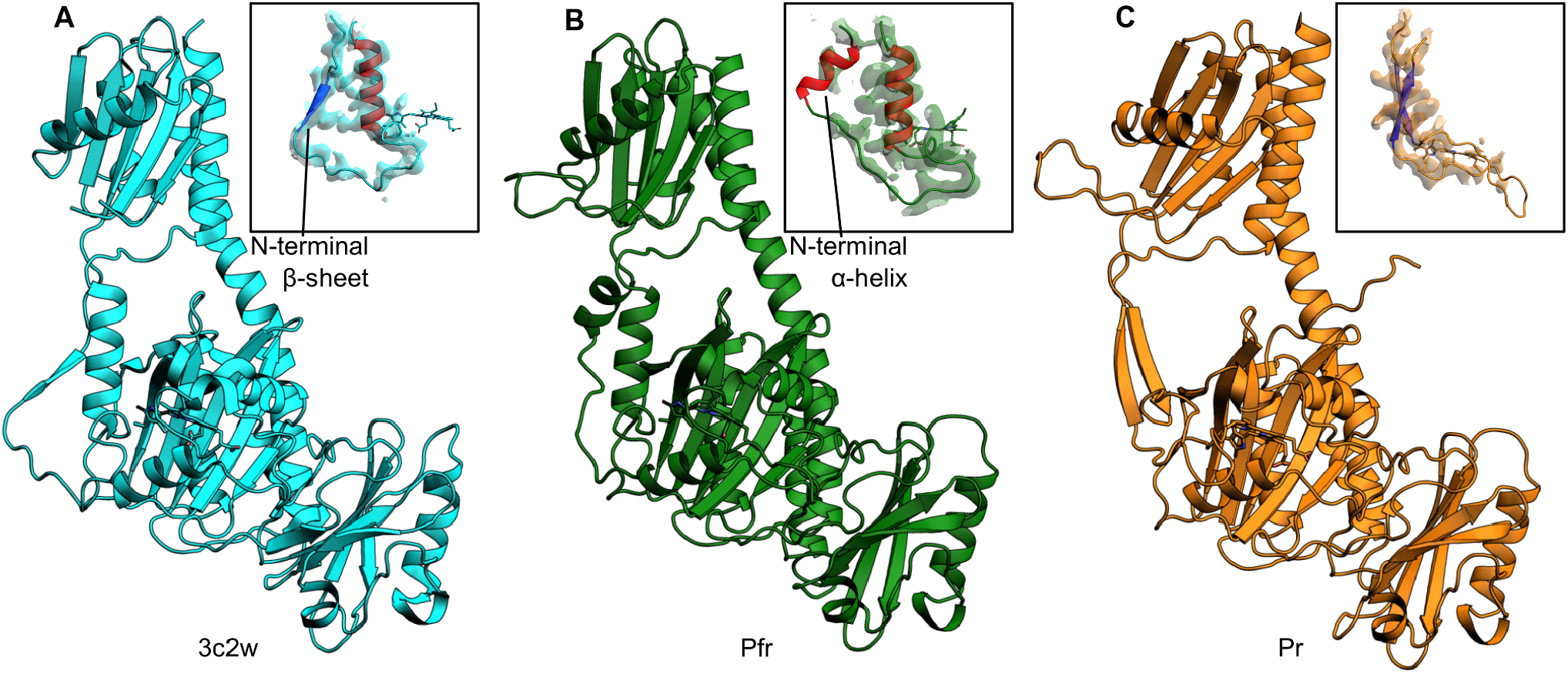
Comparison of the *Pa*BphP photosensory module in the crystal structure and the cryo EM structures. **A** Crystal structure as presented by Yang et al^8^. The tongue (inset) presents a β-sheet conformation on the N-terminal strand and an α-helical conformation on the C-terminal strand, with the 2fo-fc map overlaid. This conformation is different from the one observed in the cryo-EM structure presented here, illustrating the conformational plasticity of the region. **B** Cryo-EM structure of the protein in its Pfr state. The tongue is packed tighter, and both the N- and C-terminal strands form α-helices. Overall RMSD between the cryo-EM and crystal structures of the photosensory module is 1.08 Å. **C** Cryo-EM structure of the protein in its Pr state. The tongue forms two antiparallel β-sheets. RMSD (excluding the output module) is 1.31 Å with the Pfr cryo-EM structure and 1.69 Å with the crystal structure.

Interestingly, the Pfr structure is open at the PHY domains, with the distance of the centroids of the PHY domains increasing by approximately 25 Å compared to Pr (Fig. 4B), although the relative angle and distance of the two domains is heterogeneous (Movie S2). This separation influences the entirety of the neck region: distances between residues L487 increase from 7.3 Å in Pr to 31 Å in Pfr, and the coiled-coil structure of the neck region appears disrupted. We observe a loss of signal from this region in the consensus EM density map in Pfr (Fig. 4A), which we assign to the increased flexibility of the output domains in lieu of their dimer contact. Interestingly, we also find that the dimerisation interface at the photosensory module changes. Helix A in protomer A assumes its relaxed orientation observed in crystal structures, rearranging the GAF dimer interface and restoring symmetry of the photosensory module (Fig. 3B)^8,23^. This rearrangement has not been observed in cryo-EM or crystallographic structures of bacteriophytochromes before.

The open structure is surprising in light of the two recently solved full-length structures of Pfr-state bacteriophytochromes, which showed a closed arrangement of the photosensory module dimer^30,31^. The missing densities in this dataset could in theory be explained by compositional heterogeneity. However, the reversible nature of the photocycle and the completeness of the structure in the far-red illuminated data point towards an unresolvable conformational heterogeneity instead. The presence and ratio of closed, Pr-like particles corresponds to the residual activity observed in the dark-incubated protein (Fig. 1B) and the spectral features of this state (Fig. S1A), further supporting the interpretation that the absence of electron density is caused by conformational heterogeneity.

### The length of the neck plays a role in the structural mechanism of deactivation

Since different degrees of opening at the neck are observed for different bacteriophytochromes, we hypothesise that the constitution of the neck is responsible for the variability in this photoresponse. To investigate this, we conducted multiple sequence alignment and analysis on the three structurally characterised and over 800 additional bacterial and fungal phytochromes that feature histidine kinase output modules. In these data we searched for patterns in the length of the neck region.

To this end, we defined a connector region that included the neck but spanned from a conserved leucine in the PHY domain (L485 in *Pa*BphP) to the catalytic histidine (H513) in the DHp (Fig. 6A). The analysis reveals significant variability in this region – and hence in the neck length. The previously reported cryo-EM structures of canonical BphPs (*Dr*BphP from *D. radiodurans* and *Sa*BphP2 from *Stigmatella aurantiaca*) presented small or absent neck opening and both belong to a group with a connector length of 39 residues (Fig. 6B)^30,31^. In *Pa*BphP, the corresponding region is only 29 residues long, which is the shortest among all observed sequences. This group contains a number of other phytochromes from non-photosynthetic bacteria. Although the constitution of the dimer interface in the neck region will also play a role, the absence of 10 residues is a probable explanation to why *Pa*BphP opens up in Pfr while *Dr*BphP and *Sa*BphP2 do not^30,31^.

**Figure 6.**
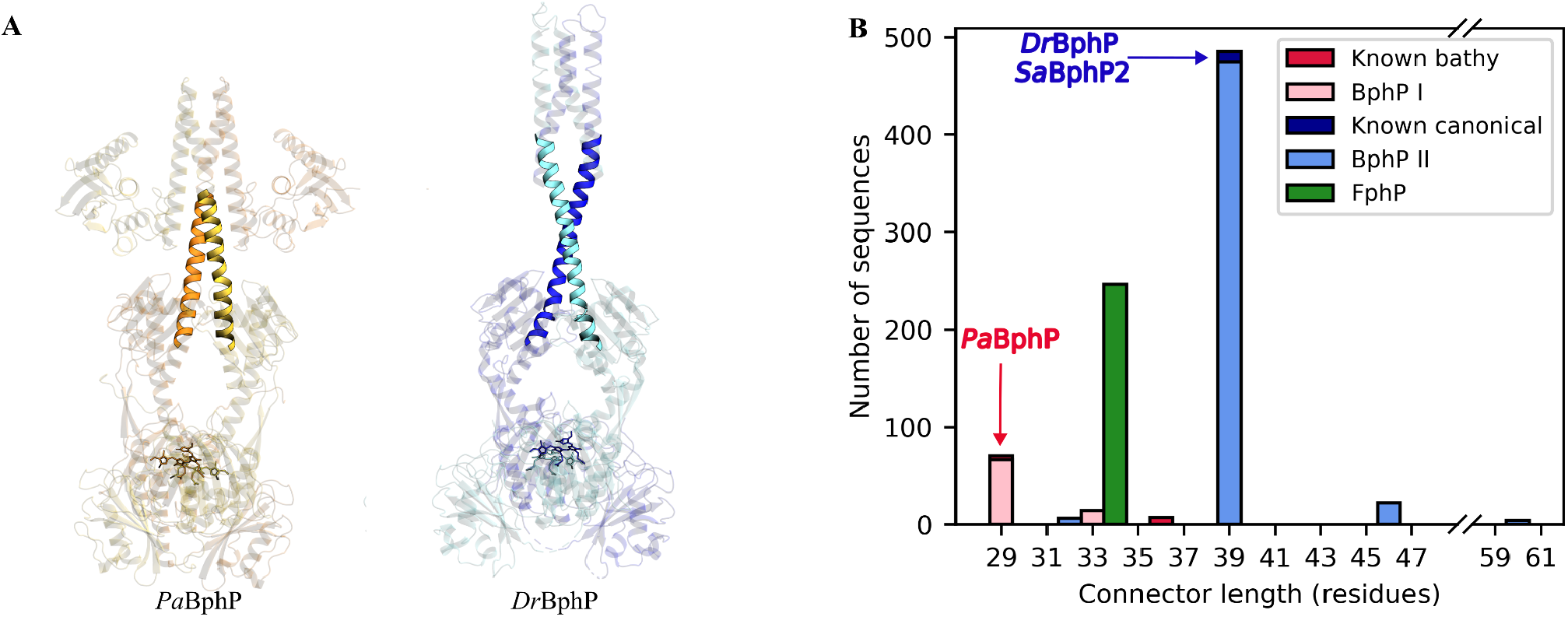
Distribution of the connector lengths across bacterial and fungal phytochrome HKs. **A** The structures of *Pa*BphP and *Dr*BphP with the connector regions highlighted^30^. **B** Histogram showing the connector lengths in the analysed sequences. Different colours represent separate clusters with connector lengths of k+n*7 residues. Proteins with known ground states are represented as darker bars, and the positions of the three bacteriophytochromes (BphP) with available full-length cryo-EM structures are indicated with arrows. Fungal phytochromes (FphP) display no variability.

### The length of the neck separates phytochromes into three clusters

We further analysed the connector length (Fig. 6) under the premise that coiled-coil linkers exhibit a full turn at a period of 7 residues, known as heptads. Given this, three separate clusters of proteins are observed, classified into separate heptad groups (l=k+n*7). Bacterial phytochromes form two clusters with l=29(/36) residues in cluster I and l=39(/32/46/60) residues in cluster II. The shift in connector lengths within either cluster corresponds to a full coil heptad, which likely leads to identical connections at the connecting loop of the DHp domain within a given cluster. The proteins with a connector length of l=33 likely belong to cluster I, with an additional half heptad, or one helical turn, of the coiled-coil. An interesting cluster is observed at l=34, which contains all analysed fungal phytochromes and forms a separate group. *Pa*BphP, along with all known bathy bacteriophytochromes, belongs to cluster I. Overall, this analysis demonstrates an important functional role for the configuration of this connector region in terms of signalling function.

## Discussion

### PaBphP functions as a kinase in its light-activated Pr state

Our kinase activity assay shows that *Pa*BphP is a functional histidine kinase. It was known that *Pa*BphP can autophosphorylate its catalytic H513 in Pr^11,36^. Here we establish that the protein transduces the phosphate signal in Pr to a response regulator protein, but that it is inactive in Pfr^11,46^. The small amount of residual enzymatic activity in dark in our assays is explained by thermal activation of the protein, which is consistent in the spectral (Fig. S1A) and structural analysis (Fig. S6). This residual activity is also consistent with available autophosphorylation data on bathy phytochromes. It is interesting to note that the on/off ratio is higher in *Pa*BphP compared to the prototypical phytochrome from *Agrobacterium fabrum* (Agp1), which was investigated by us using the same assay^12^. This supports the recent notion that bathy phytochromes are ideal for distinguishing between darkness and light. 12

The enzymatic function of *Pa*BphP is readily rationalised by our structures of the protein in Pr and Pfr. The Pr structure features a relatively stable and fully dimerised HK module, which enables the autophosphorylation reaction and phosphotransfer to the cognate response regulator. In Pfr, the dimer interface at the HK module is broken and this inhibits its kinase activity. This opening of the dimer at the PHY and DHp domains readily explains the difference of biochemical function of these states.

### Structure of PaBphP in its Pr state supports trans-autophosphorylation

Even though the histidine kinase output modules are smaller and more disordered than the photosensory modules and therefore yield consensus maps with lower resolution, the connecting loops between the DHp helices in the Pr state can for the first time be clearly traced among bacterial phytochromes. The positioning of the DHp four-helix bundle and the CA domain establishes *trans*-, but not *cis*-autophosphorylation of the catalytic histidine residue. The mobility of the domains (Supplementary movie 1) is consistent with a recent cryo EM structure of a transmembrane histidine kinase^41^. Likely, the mobility of the domains bears functional relevance, for example to maintain high entropy in the dimeric arrangement, to facilitate the autophosphorylation, or for the phosphate transfer reaction.

### Large-scale rearrangements in the quaternary structure of PaBphP during photoactivation

The degree of structural change between Pr and Pfr states in our structures of *Pa*BphP is surprising. At the HK and PHY domains, the dimer interface breaks in Pfr, and this structural shift is supported by a rearrangement of the dimerisation interface at the GAF domain. This contradicts the structural conclusions for *Pa*BphP based on modelling against time-resolved SAXS data^54^. The asymmetry in our Pr structure may indicate that the Pr state is already strained and thereby primed for opening upon photoexcitation.

The opening mechanism had been expected by extrapolation from crystal structures of bacteriophytochrome photosensory module constructs in Pr and Pfr and gel chromatography data on and gel full-length bacteriophytochromes^5,55^. However, in subsequent SAXS and cryo-EM structures of *Dr*BphP and *Sa*BphP2, this opening was not found to occur^5,30,31,55–57^. Thus, our findings finally demonstrate a mechanism of signal transduction in a full-length phytochrome where changes close to the chromophore lead to splaying apart of the PHY and catalytic DHp domains.

The structural mechanism of (de)activation of sensor histidine kinases has been vividly debated^35,58–61^. A variety of mechanisms have been proposed, but a shortage of full-length structures has made reliable assignments difficult. Here, we find that the mechanisms can indeed be very different, even in the relatively narrow group of bacterial phytochrome kinases. Figure 7 summarises the modes of signal transduction for the three full-length phytochromes solved by single-particle cryo-EM to date, which are all different. Therefore, we consider it is likely that even in the wider field of sensor histidine kinases, a variety of signal transduction mechanisms is present^39,40,62^.

**Figure 7.**
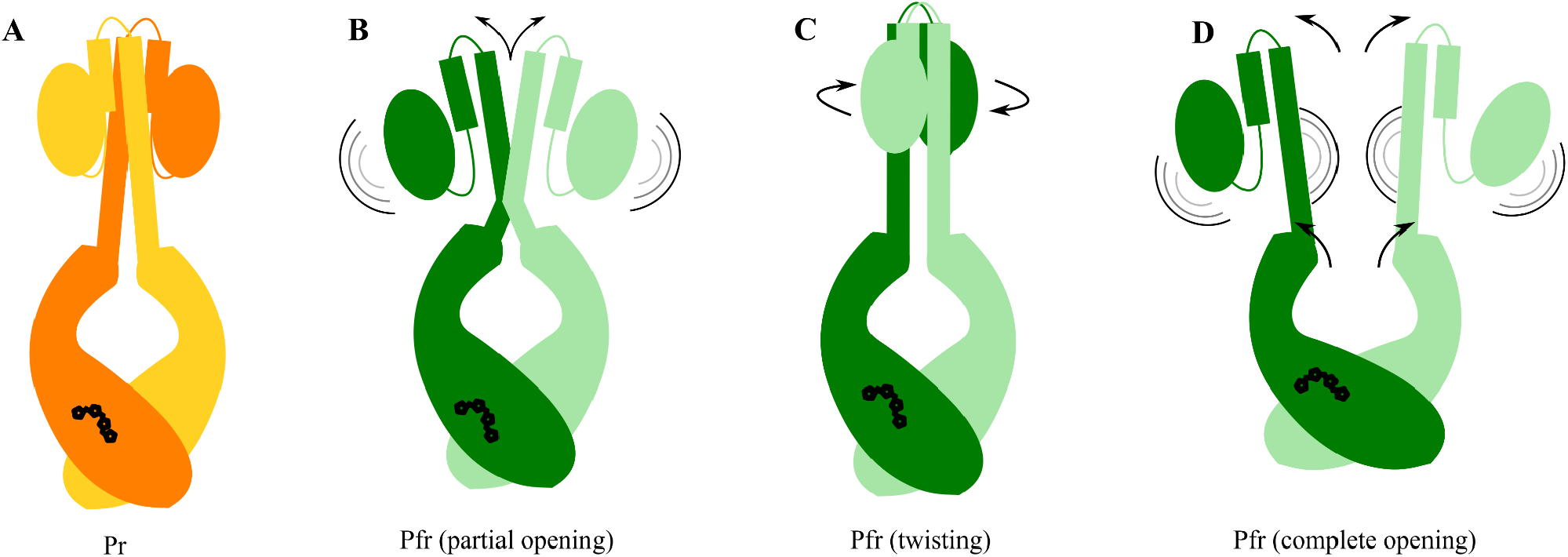
Proposed off-switching mechanisms for bacterial phytochromes. **A** Pr state, showing an ordered, symmetric output module. **B** Partially opened neck, with dynamic CA domains, as seen in Pfr-state *Dr*BphP^30^. **C** Twisted DHp domain, leading to a rearrangement of CA domain binding, as observed in Pfr-state *Sa*BphP2^31^. **D** Open photosensory module, with highly dynamic DHp and CA domains, observed for the Pfr-state *Pa*BphP in this work.

### Multiple sequence analysis of phytochromes reveals distinct clusters of connector regions

Our multiple sequence alignment analysis indicates that the variation among Pfr-state structures (Fig. 7) may be caused by the differences in the neck length. Shorter necks provide less stability for the coiled-coil structure and enable complete separation of the DHp and CA domains, resulting in more reliable off-switching of the output module. It is likely that evolution has optimised the neck length to meet these requirements. In non-photoactive bacterial sensor histidine kinases, the DHp domains are connected to coiled-coil linkers, and it is reasonable to assume that a similar evolutionary optimisation has taken place. Furthermore, this finding emphasises the importance of the length and type of the connector region when designing recombinant constructs for optogenetic applications, depending on the desired outcome of photoexcitation on the output activity.

This study presents the first comprehensive structural analysis of a bathy, full-length bacteriophytochrome, elucidating the molecular basis of its light-dependent kinase activity. We have demonstrated that *Pa*BphP operates as a kinase in its Pr form, facilitated by the ordered, dimeric arrangement of the histidine kinase module, and becomes kinase-inactive upon transitioning to the Pfr state, where the sister modules separate. Our findings reveal significant large-scale structural rearrangements between these states, highlighting a novel on-off switching mechanism that is likely to have broad implications for understanding phytochrome function across bacterial species. Additionally, our sequence analysis of the neck region across a range of bacterial and fungal phytochromes suggests an evolutionary optimisation of neck length for specific functional roles, offering new insights into the diversity and adaptability of phytochrome signalling mechanisms. This work integrates functional and structural modification of phytochromes and sets the stage for future explorations into the functional diversity, evolutionary history and optogenetic exploitation of these pivotal red light-sensing proteins.

### Experimental procedures

#### Protein preparation and purification

The plasmid used for the recombinant expression of the full-length phytochrome from *P. aeruginosa* consists of the *Pa*BphP insert (GenBank: AAG07504.1) flanked by a C-terminal His6-tag, and the pET21b(+) plasmid backbone (Novagen). The heme oxygenase plasmid used to provide sufficient biliverdin incorporation during protein expression was kindly provided by Prof. Janne A. Ihalainen. The expression and purification of *Pa*BphP were carried out following the procedure outlined below. Double-transformed *Escherichia coli* strain BL21 (DE3) cells were cultured in LB media supplemented with 50 μg/ml kanamycin at 37°C until *OD*_600_ reached value 0.6. 1 mM IPTG and 5-aminolevulinic acid were added to induce expression of *Pa*BphP and heme oxygenase, and to aid biliverdin synthesis, respectively. Protein expression was done overnight at 18 °C. Following cell lysis in lysis buffer (50 mM Tris, 150 mM NaCl, 10% glycerol, 10 U/ml of DNase 1 and one tablet of cOmplete EDTA-free protease inhibitor (Roche) at pH 8.0) using an EmulsiFlex C3 homogeniser (Avestin), the lysate was briefly incubated with a molar excess of biliverdin hydrochloride. The His6-tagged protein was then purified using Ni-NTA affinity chromatography (HisTrap™ HP 5 ml, Cytiva) by washing the bound sample with wash buffer (50 mM Tris and 1 M NaCl at pH 8.0) and eluting with elution buffer (50 mM Tris, 50 mM NaCl, 300 mM imidazole at pH 8.0); followed by size-exclusion chromatography (HiLoad™ 16/600 Superdex™ 200 pg, Cytiva) in SEC buffer (30 mM Tris at pH 8.0). The purified protein was collected at a concentration of 3 mg/ml and flash-frozen for storage.

#### Spectroscopy

The spectroscopic measurements were conducted using AvaSpec-ULS2048CL-EVO an (Avantes) spectrometer in combination with an Avalight-XE-HP (Avantes) light source, in a quartz cuvette with 10 mm light path. The protein sample was kept in complete darkness for 24 hours for the dark state spectrum. For the far-red illuminated spectrum, the sample was illuminated with a 780-nm LED light source with a total energy of 1 mJ/mm^2^. Dark reversion time was measured at room temperature, by recording spectra of the protein in dark at regular intervals immediately after far-red illumination. The degree of Pr content was determined by the absorption intensity at 700 nm.

#### Phos-tag assay

As the cognate response regulator of *Pa*BphP is not known, the Phos-tag assay was conducted with the response regulator from *D. radiodurans* (*Dr*RR). To generate phosphorylated *Dr*RR (p-*Dr*RR), *Dr*RR was pre-treated with 50 mM acetyl phosphate, as described previously ^30^. Both kinase and phosphatase reactions were carried out in a buffer solution (25 mM Tris/HCl pH 7.8, 5 mM MgCl_2_, 4 mM 2-mercaptoethanol, 5% ethylene glycol) with *Pa*BphP and *Dr*RR/p-*Dr*RR (2–4 µg each). The reactions were initiated by adding 1 mM ATP to the mixture and incubated at 25 °C, either in the dark or under saturating far-red (785 nm) light, for 20–30 minutes. The reactions were halted by adding 5*SDS-PAGE loading buffer. For the detection of p-*Dr*RR amount via mobility shift, we employed the Zn^2+^-Phos-tag® SDS-PAGE assay (Wako Chemicals) according to manufacturer’s instructions. The samples were run with 40 mA in 9% SDS-PAGE gels containing 50 µM Phos-tag acrylamide at room temperature.

#### Sample preparation for cryo-electron microscopy

Prior to grid preparation, the protein buffer was exchanged to a solution containing 80 mM Tris, 10 mM MgCl_2_ and 150 mM CH_3_CO_2_K with a pH of 7.8, and the protein concentration was set to 1.5 mg/ml. The protein was handled under dim 530-nm illumination during the preparation of dark Pfr state samples. For Pr state samples, an additional illumination step with a 780 nm light source (1 mJ/mm^2^) was performed immediately before applying the solution on the grids (<10 s before freezing).

UltrAuFoil (Quantifoil) R 1.2/1.3 (300 mesh) grids were glow discharged for 60 s in an easiGlow (Pelco) glow discharge cleaning system before applying 3 μl protein solution and freezing in liquid ethane using a Vitrobot mark IV (Thermo Scientific), set to 100% humidity and 4°C.

#### Cryo-electron microscopy data collection

The prepared grids were loaded in a Titan Krios G2 (Thermo Scientific) transmission electron microscope equipped with a K3 (Gatan) detector and a 20 eV BioQuantum (Gatan) energy filter, operated at 300 keV. Data were collected with the following settings: 0.828 Å/pixel (105kx magnification), 50 e^-^/Å^2^ total electron dose, -0.8 - -2.0 μm defocus range. Automated collection was performed using the EPU software (Thermo Scientific). A total of 30,336 movies were collected for the Pr state, and 15,496 movies for the Pfr state.

#### Cryo-electron microscopy data processing

The micrographs were processed using Cryosparc v4.2.1 (Structura Biotechnology). The movies were pre-processed by patch motion correction and patch CTF estimation, and low quality micrographs were discarded based on CTF fit, relative ice thickness and average intensity. Initial particle picking was done using blob picking for both datasets.

In the Pr dataset, different views of the closed conformation were used for template picking, and a Topaz model was trained on a clean particle set for the final round of picking. After *ab initio* reconstruction and heterogeneous refinement, a set of 889,461 particles was used for local refinement of the photosensory module. 3D variability analysis was used to uncover the structural heterogeneity. Further 3D classification of the particle set resulted in a subset of 179,058 particles, which were used to reconstruct the output module using non uniform refinement. An initial molecular model was aligned to the two structures, and the Phenix function ‘Combine focused maps’ was used to generate a composite map.

In the Pfr dataset, initial reconstructions were built for both the open and the closed conformations, and separate Topaz models were trained to pick particle sets for either conformation. The low number of total particles has prevented the reconstruction of a high resolution closed conformation structure. For the open state, *ab initio* reconstruction and heterogeneous refinement has resulted in a set of 1,103,315 particles. 3D variability analysis was used to uncover the structural heterogeneity. Further 3D classification has resulted in a subset of 239,820 particles, which were used for the final reconstruction using non-uniform refinement followed by local refinement.

#### Model building and refinement

Models for both the Pr and Pfr state were built using the ISOLDE (v1.5) extension of ChimeraX (v1.5)^63^. An initial model was generated using ColabFold (v1.5.5), and the protomers were separately fitted into the density maps as rigid bodies^64^. To add biliverdin to the model, the GAFF force field in combination with the AM1-BCC charge model was used to generate the molecule LBV in antechamber^65–67^. The finished models were refined against the electron density maps using the Phenix (v1.20) function ‘Real-space refinement’.

#### Multiple sequence analysis

BLAST was used on six different phytochrome sequences with the following Uniprot accession codes: *Dr*BphP (*D. radiodurans* - Q9RZA4), *Pa*BphP (*P. aeruginosa* - Q9HWR3), *Sa*BphP2 (*S. aurantiaca* - Q097N3), *Af*BphP (*A. fabrum* - Q7CY45), *En*FphA (*Emericella nidulans* - Q5K039) and Cph1 (*Syneochocystis sp*. PCC6803 - Q55168)^68^. The E-Threshold was set to 0.1, the BLOSUM62 matrix was used and 250 hits per sequence with a minimum sequence identity of 30% were chosen^69^. Double hits were removed, which led to a total sequence number of 859 from 361 different families. The sequences were then aligned to the catalytic histidine in the output module.

Multiple sequence alignment (MSA) was performed using the Clustal Omega web server^70^. CLUSTAL W was used to generate the MSA^71^. Afterwards, trimAl was used to cut the sequences at two reference positions, corresponding to residues L485 (within the PHY domain) and H513 (within the DHp domain) of *Pa*BphP^72^. The residues spanning the two reference positions were then counted and plotted as a histogram (Fig. 6B), using an in-house python script.

## Acknowledgements

We acknowledge the use of the Cryo-EM Uppsala facility for cryo-EM specimen preparation and screening, funded by the Department of Cell and Molecular Biology, the Disciplinary Domains of Science and Technology and of Medicine and Pharmacy at Uppsala University. We thank Daniel Larsson and Anna Sundborger-Lunna for their assistance. We thank Dr. Elina Multamäki (University of Helsinki) for her help with conducting the *Pa*BphP phosphorylation assays.

The cryo-EM data was collected at the Cryo-EM Swedish National Facility funded by the Knut and Alice Wallenberg, Family Erling Persson and Kempe Foundations, SciLifeLab, Stockholm University and Umeå University. We are grateful for the help provided by Marta Carroni, Mathieu Coincon, Julian Conrad, Dustin Morado and Karin Walldén during data collection.

Parts of the data processing were enabled by the supercomputing resource Berzelius provided by National Supercomputer Centre at Linköping University and the Knut and Alice Wallenberg foundation.

SW acknowledges funding from the Swedish Research Council (grant number 2021-05101). HT acknowledges the Research Council of Finland (grant 330678) for funding.

## Author contributions

SW designed the project, SB, HT, SW planned the project and the experiments, SB and PM purified the protein, SB produced the grids and performed the single-particle cryo EM data acquisition and refinement and interpreted the structures together with SW, HT performed the phosphorylation assays, LG performed the multiple sequence alignment, SW and HT secured funding, SW supervised the project, SB, PM and SW wrote the paper with input from all authors.

## Data availability

The data that support this study are available from the corresponding authors upon request. Raw microscopy images have been deposited in EMPIAR (EMPIAR-12093). EM density maps are available in EMDB: EMD-19989 (Pfr), EMD-19981 (Pr composite map), EMD-50007 (Pr consensus refinement), EMD-50008 (Pr photosensory module local refinement), EMD-50009 (Pr output module local refinement). PDB codes for the reconstructed models are 9EUY (Pfr) and 9EUT (Pr).

## Supplementary information

Document S1 containing Figures S1 to S6; Movies S1 to S2 as separate files.

**Movie S1 (separate file)**. Structural heterogeneity in the Pr particle set revealed by 3D variability analysis (Cryosparc). While the photosensory module acts as a rigid structure, the neck swings considerably, causing blurring of the output module in global reconstructions.

**Movie S2 (separate file)**. Structural heterogeneity in the Pfr particle set revealed by 3D variability analysis (Cryosparc). The dimeric interface at the GAF domain is rigid, but the PHY domains of the protomers move relative to each other, limiting resolution. The output module is not resolvable due to its small size and large freedom of motion.

