## Supplementary figures and tables for "Cryo-EM structures of a bathy phytochrome histidine kinase reveal a unique light-dependent activation mechanism"

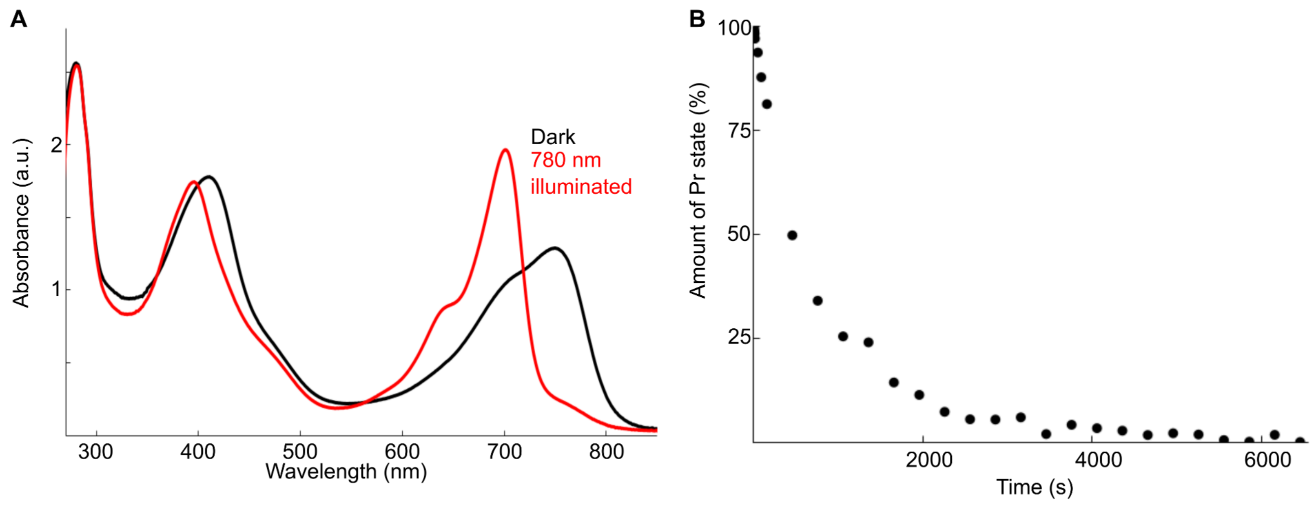


**Figure S1. Spectral characteristics of *Pa*BphP. A** The UV-Vis absorption spectrum of *Pa*BphP in its dark-adapted state (black line) and immediately after illumination with saturating 780 nm light (red line). The absorption at 400 nm is high compared to other phytochromes and there is an unusual shoulder present at 475 nm. We tentatively assign this absorption to degraded excess biliverdin present in the sample buffer, but not observed during the cryo EM data refinement process. **B** Dark reversion of the protein from the Pr state at room temperature. The amount of Pr state was determined as a function of $Abs_{700}$.The time scale for dark reversion allows capturing the Pr state by conventional cryo-EM specimen preparation methods.

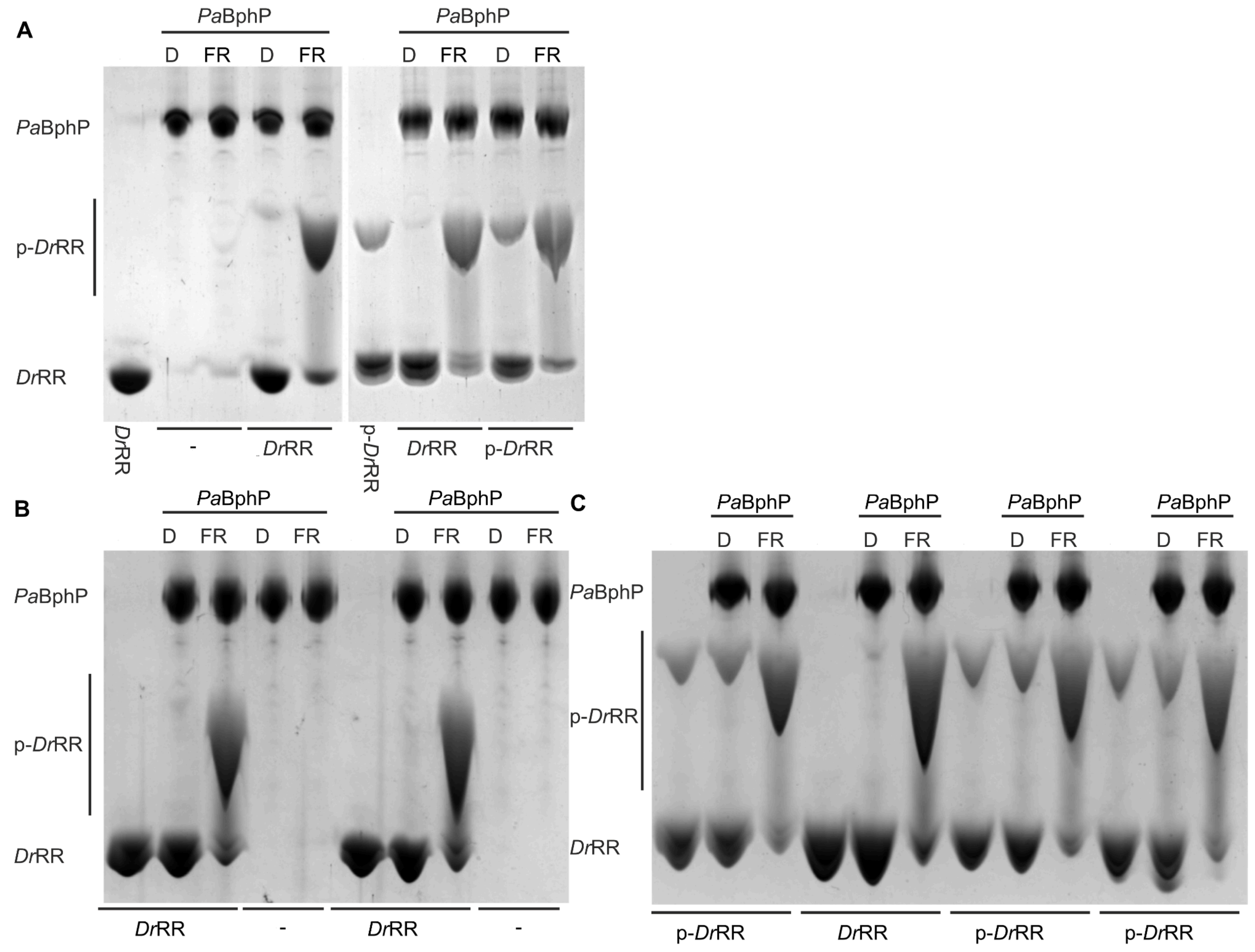


**Figure S2. Extended gel images of the *Pa*BphP phosphorylation assay. A** The native bands of *Dr*RR and p-*Dr*RR without the addition of *Pa*BphP; the effects of *Pa*BphP on the *Dr*RR - p-*Dr*RR equilibrium when incubated with *Dr*RR or p-*Dr*RR in the dark or under far-red illumination. **B-C** Repeats of the assay to confirm the results of panel a and Figure 2 in the main text

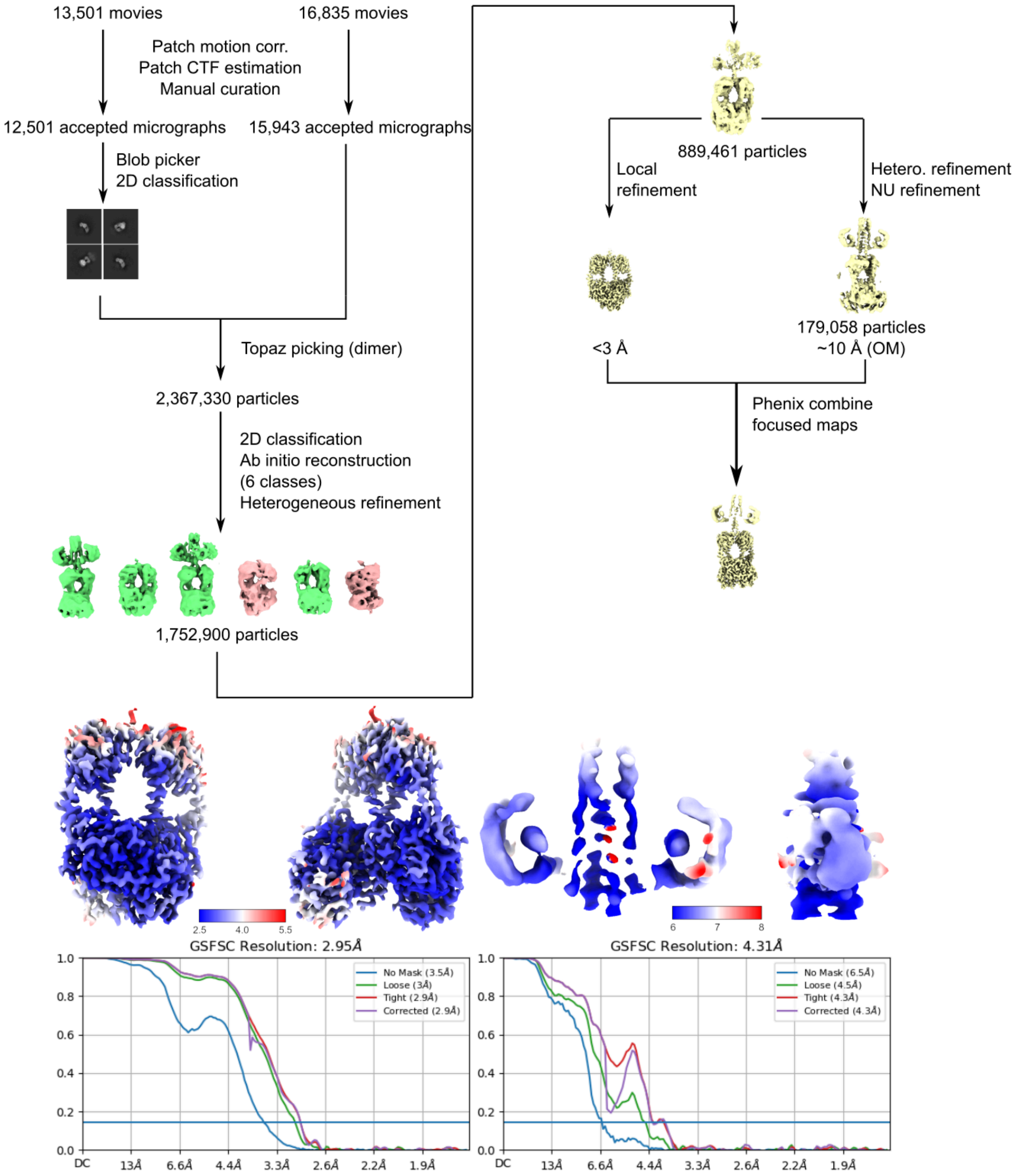


**Figure S3. Processing of the far-red illuminated cryo-EM data.** A total of 30,336 exposures were collected, of which 28,444 were selected after filtering based on CTF fit, average ice thickness and total in-frame motion. Initial particle assessment was done by blob picking, and the final particle set was picked using the Topaz deep picking algorithm. Junk particles were discarded during 2D classification and heterogeneous refinement, and a consensus refinement was created using 889,461 particles. To achieve higher resolution, the photosensory and output modules were refined separately, and the separate volumes were combined into a composite map.

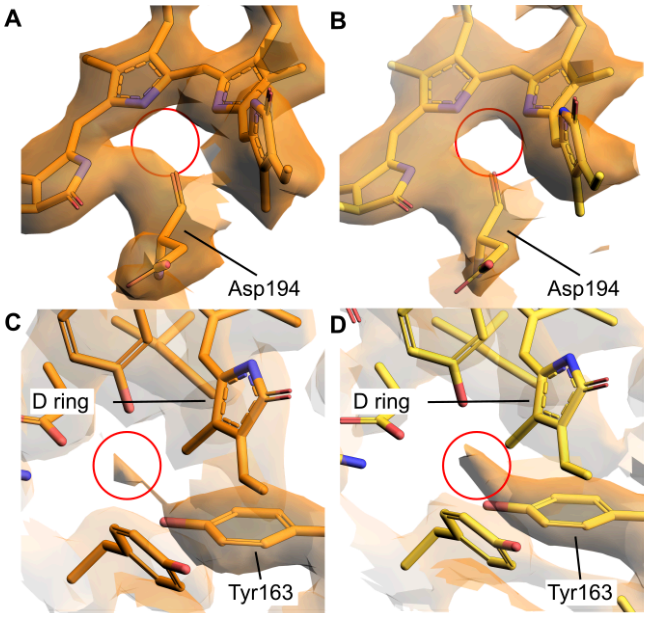


**Figure S4. Presence of structural waters in the density.** The pyrrole water is not observed either in protomer A (**A**) or in protomer B (**B**). An ordered water is present between the biliverdin D-ring and residue Y163 in protomer A (**C**) and protomer B (**D**) of *Pa*BphP in the Pr state (not modelled).

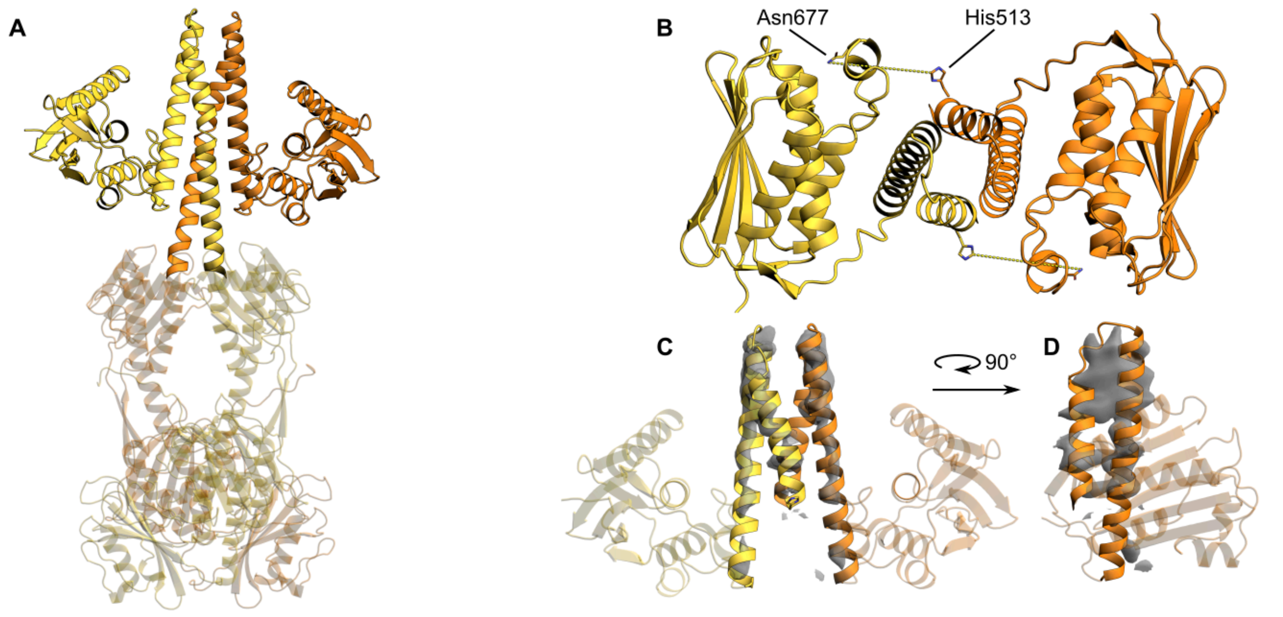


**Figure S5. Configuration of the output module of *Pa*BphP.** The direction of the DHp connecting loop allows *trans*- but not *cis*-autophosphorylation of the catalytic histidine residue. **A** The output module of *Pa*BphP. **B** Top view of the output module, with the side chains of the catalytic histidine and the ATP-binding asparagine highlighted. The distance between H513 of protomer A and N677 of protomer B (and *vice versa*) is measured to be approximately 16 Å, although at the achieved resolution this measurement is of low certainty. **C-D** The density maps around the DHp domain from front (**C**) and side (**D**, only protomer A shown), supporting the *trans* configuration of the histidine kinase.

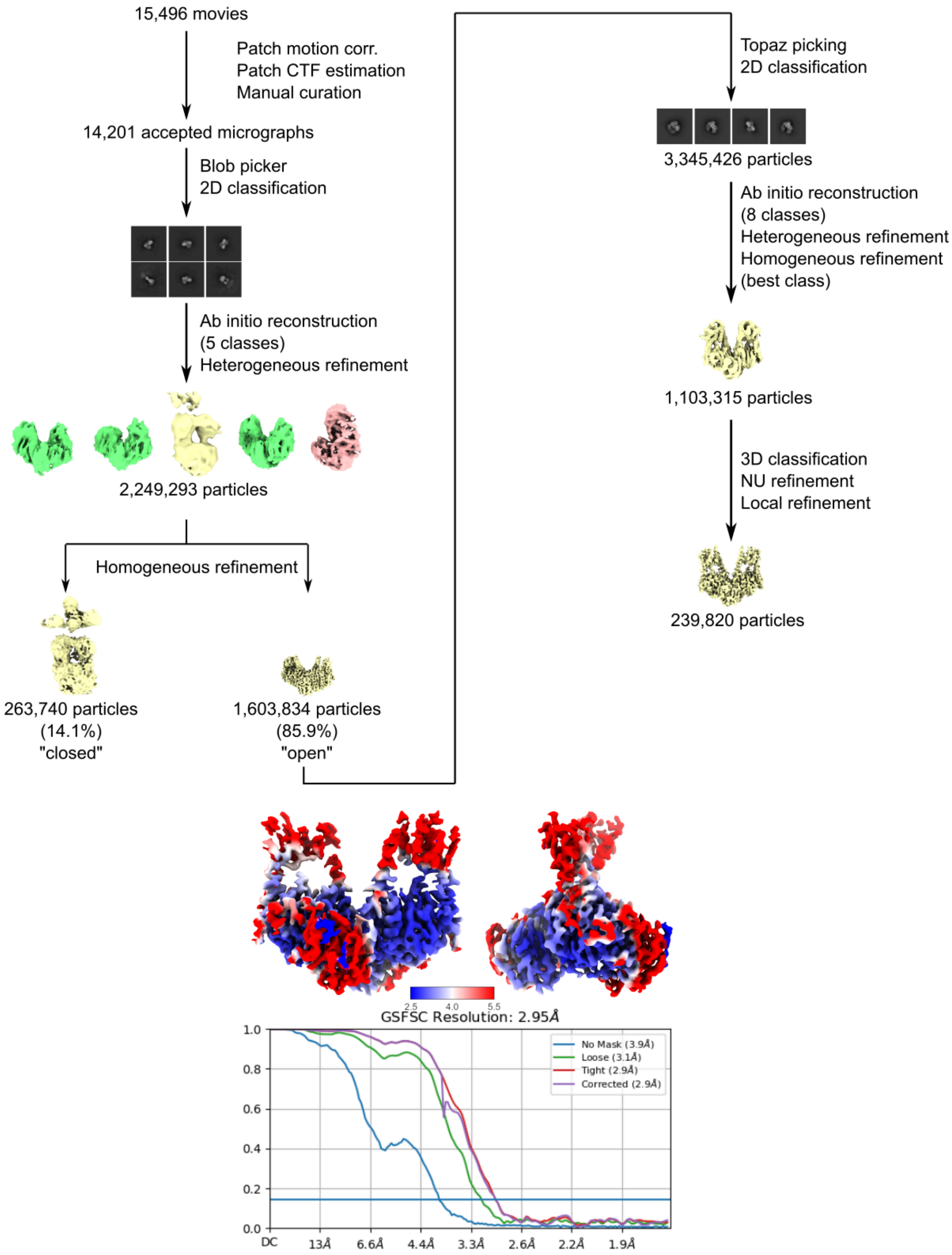


**Figure S6. Processing of the dark cryo-EM data.** A total of 15,496 exposures were collected, of which 14,201 were selected after filtering based on CTF fit, average ice thickness, and total in-frame motion. Initial particle assessment was done by blob picking, and the final particle set was picked using the Topaz deep picking algorithm. Junk particles were discarded during 2D classification and heterogeneous refinement, and two separate conformations were reconstructed from the data, labelled "closed" and "open", based on the relative positioning of the photosensory modules. Further refinement of the closed conformation was not successful due to the low amount of real particles. The final refinement of the open conformation was done using 239,820 particles. The local resolution map shows that the resolution decreases in the PHY domains compared to the PAS/GAF domains, which we assign to the flexibility of the dimer (see also Movie S2). Regardless, the backbone can be reliably traced throughout the entire photosensory module.

| | | Pr | Pfr |
|---|---|---|---|
| **Data collection** | | | |
| | Voltage (keV) | 300 | 300 |
| | Magnification | 105,000x | 105,000x |
| | Pixel size (Å/px) | 0.828 | 0.828 |
| | Defocus range (um) | -0.8 - -2.0 | -0.8 - -2.0 |
| | Electron dose (e/Å2) | 50 | 50 |
| **Reconstruction** | | | |
| | Accepted micrographs | 30,336 | 15,496 |
| | Imposed symmetry | C1 | C1 |
| | Particles (consensus refinement) | 889,461 | 1,103,315 |
| | Nominal resolution (Å) | 3.79 | 3.37 |
| | Particles (PSM local refinement) | 889,461 | 239,820 |
| | Nominal resolution (Å) | 2.95 | 2.95 |
| | Particles (HK local refinement) | 179,058 | - |
| | Nominal resolution (Å) | 4.31 | - |
| **Model building** | | | |
| | Initial model | AlphaFold2 | AlphaFold2 |
| | Non-H atoms | 11,456 | 7846 |
| | Residues | 1456 | 984 |
| | Ligands | 2 | 2 |
| | B factor (protein) | 75.48 | 81.09 |
| | B factor (ligand) | 14.54 | 35.87 |
| | Bond length/angle RMSZ | 0.31/0.58 | 0.27/0.57 |
| | Molprobity score | 1.19 | 1.10 |
| | Clashscore | 3 | 3 |
| | Ramachandran outliers | 0 | 0 |
| | Sidechain outliers | 1.00% | 0.40% |
| | Q-score | 0.402 | 0.466 |

**Table S1. Data collection, processing and refinement statistics of the Pr and Pfr datasets.**
